# Synergistic positive feedback underlying seizure initiation

**DOI:** 10.1101/2021.02.28.433224

**Authors:** Robert T. Graham, R. Ryley Parrish, Laura Alberio, Emily L. Johnson, Laura J. Owens, Andrew J. Trevelyan

## Abstract

Seizure onset is a critically important brain state transition that has proved very difficult to predict accurately from recordings of brain activity. Here we show that an intermittent, optogenetic, stimulation paradigm reveals a latent change in dendritic excitability that is tightly correlated to the onset of seizure activity. Our data show how the precipitous nature of the transition can be understood in terms of multiple, synergistic positive feedback mechanisms: raised intracellular Cl^-^ and extracellular K^+^, coupled to a reduced threshold for dendritic plateau potentials, and which in turn leads to a switch to pyramidal burst firing. Notably, the stimulation paradigm also delays the evolving epileptic activity, meaning that not only can one monitor seizure risk safely, it may even have an additional anti-epileptic benefit.

**One Sentence Summary:** Rapid transitions into seizures arise from mutually accelerating feedback loops, involving changes in dendritic excitability

## Main Text

Brain state transitions are fundamental for neocortical function. They are readily apparent from passive observation of brain rhythms(Buzsáki, 2006), but can be difficult to predict from such recordings, suggestive of two things: that the underlying cause is cryptic to the recording methods being used, and that the binary switch involves mechanisms that accelerate the state transition and stabilize the new state. This issue has major clinical ramifications in the field of epilepsy, because the inherently unpredictable nature of seizures is one of the biggest problems faced by people living with this condition.

Various studies have framed brain state transitions in terms of attractor states, in which positive and negative feedback mechanisms are involved in pushing the system away from the phase space bifurcation, into one, or other, attractor. The action potential threshold exemplifies such a bifurcation point, where depolarizing conductances, that are themselves activated by depolarization, constitute a strong positive feedback that accelerates the transition once the threshold is surpassed (in this instance, the threshold is the bifurcation point), but below which, the membrane returns to its resting potential, the system’s stable point.

Action potentials involving NMDA receptors and voltage-gated Ca^2+^ channels (VGCCs) – so called “dendritic spikes” or “plateau potentials”(Schiller et al., 2000)-were long ago postulated to underlie epileptic discharges, in an influential model of the paroxysmal depolarizing shift(Johnston and Brown, 1981; Traub and Miles, 1991; Traub et al., 1989; Wong and Prince, 1979), but their involvement in the actual ictogenic transition has received little attention. Instead, investigations into seizure initiation, in recent years, has focused almost overwhelmingly upon interneuronal function (Avoli et al., 2016; Chang et al., 2018a; Khoshkhoo et al., 2017; Parrish et al., 2019; Trevelyan and Schevon, 2013), chloride homeostasis (Alfonsa et al., 2016; Alfonsa et al., 2015; Burman et al., 2019; Cossart et al., 2005; Dzhala et al., 2010; Pallud et al., 2014; Pavlov et al., 2013) and extracellular potassium levels (Somjen, 2004; Somjen and Giacchino, 1985). On the other hand, multiple laboratories have demonstrated changes in expression of various ion channels that influence dendritic excitability, including voltage-gated Ca^2+^ channels(Su et al., 2002), I_Nap_(Kearney et al., 2001), I_A_(Bernard et al., 2004), I_h_(McClelland et al., 2011; Santoro and Shah, 2020) and SK-type K^+^ channels(Cai et al., 2007), that are associated with epileptic phenotypes (see also reviews (Poolos and Johnston, 2012; Tang and Thompson, 2012)). Notably, dendritic plateau potentials have been implicated in changes in perceptual thresholds (Takahashi et al., 2016), suggestive of their involvement in other transitions of cortical network function. We now present data showing that dendritic excitability is a prime determinant of seizure initiation which links the various synergistic positive feedback mechanisms, and that together, propel the brain from normal function into a seizure state.

## Proactive assays reveal binary switch in network excitability before seizure onset

We took a combined optogenetic, electrophysiological, and imaging approach, using various acute seizure models, to identify key changes in cortical networks that were most closely associated with the initiation of seizure activity. Pharmacologically induced seizure-like activity (ictal events) develops with a highly characteristic pattern and time course (Figs 1, S1), and once started, the seizure-like activity (SLA) tends to persist. This stereotyped evolving pattern is indicative of an ictogenic pacemaker, driving the network persistently towards the seizure state. The utility of these models is that they represent network systems that progress through the entire spectrum, from normal to epileptic, within a circumscribed time period, thus allowing us to identify the moment when the network crosses some critical threshold. One can then investigate how the time course of change for other network parameters aligns to this threshold point, along the ictogenic pacemaker ramp.

**Fig. 1.**
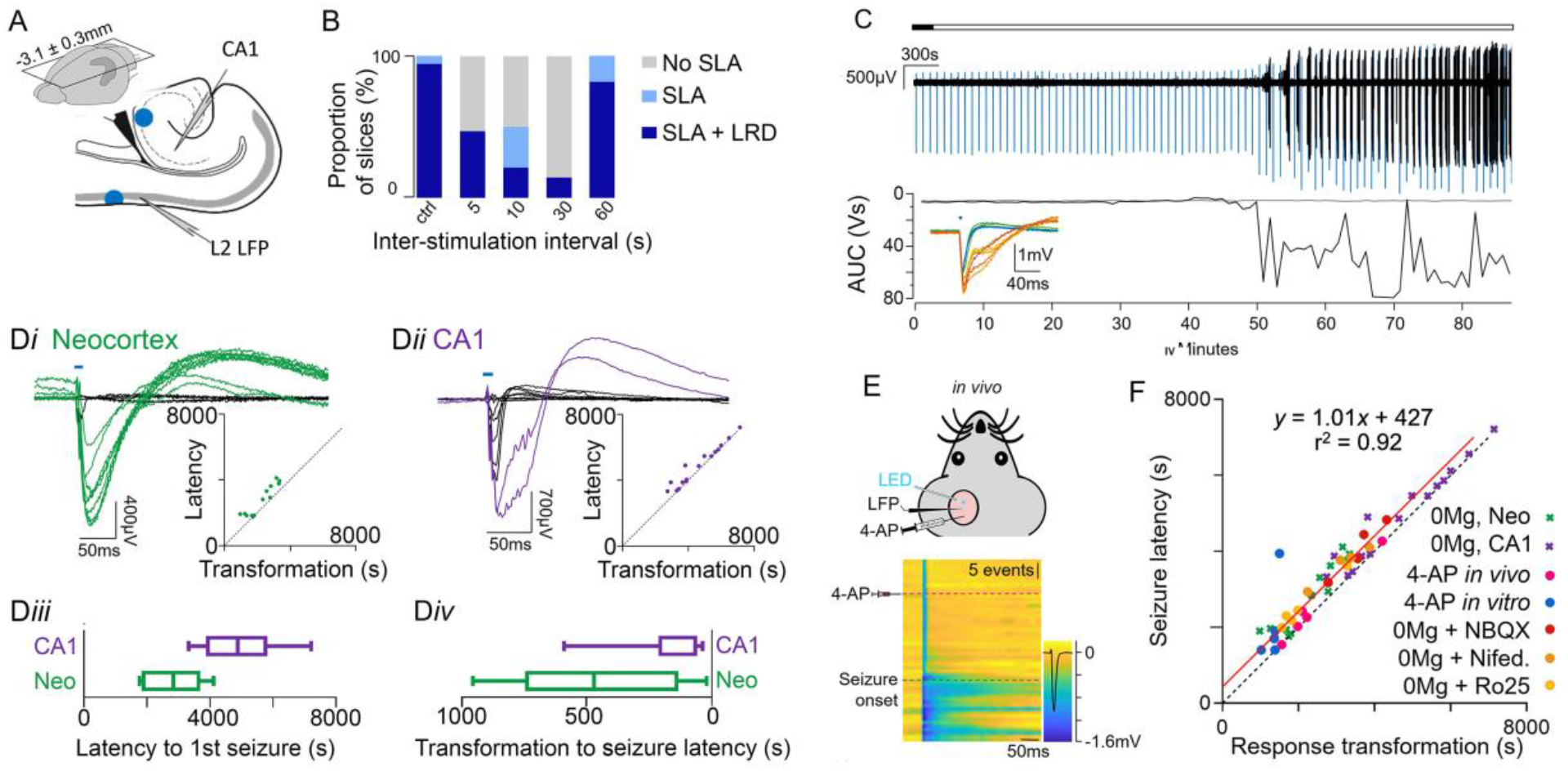
All-or-nothing transformation of response to optogenetic activation of pyramidal cells occurs shortly before seizure onset. (A) Schematic showing the *in vitro* recording arrangement in horizontal brain slices prepared from young adult mice expressing channelrhodopsin in pyramidal cells. (B) The effect of intermittent optogenetic stimulation at 4 different stimulation frequencies (every 5s, 10s, 30s and every minute) relative to unstimulated slices (ctrl). Dark blue shows the number of slices that progressed all the way to the end-stage activity pattern (“late recurrent discharges”, LRD) indicative of hippocampal involvement, while the light green indicates how many slices showed neocortical seizure-like activity (SLA). Note that all slices reaching the LRD phase show prior SLA. The recordings all lasted 1 hour after washing out Mg^2+^. Grey indicates those slices that showed no SLA in this time. (Ci) Illustrative recording of neocortical activity. 500ms epochs around each stimulus are coloured blue, to distinguish the spontaneous activity (black) from the evoked activity (blue). (Cii) Two consecutive evoked response at the time of the fEPSP transformation, that happened just before the first seizure-like event. The lower panel shows, on the same time scale, the change in amplitude and half-width, relative to baseline events. (D) Examples of the transitions shown in the neocortical (green) and CA1 (purple) evoked responses (baseline events subtracted). The difference in latency to first SLA in both areas is shown in Diii, and the latency from the transformation of the evoked EPSP to the first SLE is shown in Div. (E) Schematic showing the *in vivo* recording arrangement and the stepwise change in the evoked response. (F) Summary plot of all experiments, showing the close correlation between the time of the fEPSP transformation (abscissa) and the time of the first seizure (ordinate). The red line is the linear fit to the data (gradient = 1.0055 ± 0.0403; no significant difference from the line of equality (black), gradient = 1; p = 0.5517).

We first examined ictogenesis in brain slices, bathed in artificial CSF (aCSF, Fig 1A-D) either lacking Mg^2+^ ions (0Mg^2+^), or supplemented with the convulsant, 4-aminopyridine (4-AP, Fig S1). Notably, direct pharmacological effect occurs quickly, as assayed by the change in the postsynaptic response (normalized increase in the field excitatory postsynaptic potential, fEPSP = 0.12 ±0.08; n = 4) to a short optogenetic activation of pyramidal cells (495 ± 75s, n = 4; p<0.001, Welch’s t-test; Figs S2, S3), whereas the latency to the first seizure-like activity is almost an order of magnitude longer (in 0Mg^2+^, latency = 2840 ±1159s; in 100µM 4-AP, latency = 2068 ±1067s, n = 13 and 5 respectively, Fig S2). This suggests that the induced epileptic activity arises through secondary, and as yet undetermined, network changes induced by the experimental manipulation. We therefore set out to identify which cellular phenomena showed changes that coincided with the onset of seizure-like events (SLE). Since the various models differ in their ictogenic mechanism (Codadu et al., 2019a), we asked whether they shared common features at the moment of SLE onset.

Optogenetic stimulation was delivered to superficial layers (centred on L1) of neocortex, and the response was recorded from an extracellular electrode positioned in layer 2/3, laterally displaced and outside the direct site of illumination (Fig 1A). Intermittent stimulation, every 5s, 10s, or 30s, all markedly reduced the rate of epileptiform evolution in this model (Fig 1B) (Chiang et al., 2014; Ladas et al., 2015). While such an anti-epileptic effect would ordinarily be considered beneficial, in this instance we were interested in examining the transition into a seizure, so increased the stimulation interval to 1 min, which, while providing a lower resolution assay, allowed seizure activity to start in a timely fashion (Fig 1A, S4). In normal aCSF, the response to optogenetic stimulation remained stable for as long as the experiment lasted (Fig S4B). On washing out Mg^2+^ ions, after a rapid initial change attributable to the removal of the voltage-dependent blockade of NMDA receptors (NMDA-Rs), the fEPSP remained stable for many minutes, until there occurred a further sharp increase in both amplitude (normalised mean = 1.9 ±0.4) and duration (normalised mean = 5.04 ±2.38; latency range = 1900-4050s; Fig 1C,D). This transformation often happened within a single stimulation period (1min), and thereafter persisted, indicative of a binary, all-or nothing change. Shortly after this transformation, in all brain slices, the first sustained SLE occurred (406.8 ±316.3s; range of <1 – 12mins; n = 16; correlation of latencies of fEPSP transformation and first SLE, r^2^ = 0.86, p < 0.01).

The low Mg^2+^ model has an interesting feature that hippocampal epileptiform discharges start much later (4879.1 ±1222.8s; n = 16) than those in neocortex (Figs 1Diii, S5) (Codadu et al., 2019b). We therefore examined how the fEPSP transformation occurred in the two brain areas, alternating stimulation of neocortex and the CA3 Schaffer collateral pathway (Fig S3) to avoid crosstalk or interference between responses. Even though both brain areas experienced the removal of Mg^2+^ ions simultaneously, the transformation of fEPSPs at the two sites did not coincide; rather, the hippocampal fEPSP occurred long after both the neocortical transformation and the initial neocortical seizure activity (Fig 1Diii), but again immediately prior to the onset of local hippocampal seizure activity (Fig 1Div). In both cases, the fEPSP transformation maintained its close temporal association, immediately prior to local recruitment to epileptiform activity (hippocampal fEPSP / SLE latency correlation, r = 0.94, p < 0.01, n = 16). We additionally used a variety of pharmacological blockers to alter the rate of epileptiform evolution, and in all cases, the strong temporal association between the fEPSP transformation and the start of seizure activity remained (Fig 1F). Notably, the same temporal association was also seen in slices bathed in 4-aminopyridine, in which epileptiform activity evolves by a different mechanism, primarily by destabilizing interneuronal activity. Finally, the transformation of the fEPSP was also apparent, immediately before seizure onset, in an acute *in vivo* model, induced in urethane-anaesthetized adult mice by a focal injection of 4AP (n=5) (Fig 1E).

Collectively, these data showed a precise, one-to-one relationship between the transformed fEPSP and the onset of seizure-like activity (Fig 1F, r^2^ = 0.924, gradient = 1.0055 ± 0.0403; not significantly different from 1, 2-tailed t-test, p = 0.5517), and was robust across a wide range of experimental manipulations that alter the rate of acutely evolving epileptiform activity, both *in vitro* and *in vivo*. In every instance, the fEPSP occurred prior to the time of the first seizure-like event (Fig. 1F; y intercept = 427 ± 151s; 1-tail t-test significantly different from 0, p < 0.001). We conclude therefore that using intermittent stimulation, to proactively assay the network excitability, provides a very direct predictor of when seizure-like activity starts in acute pharmacological models.

## The transformed fEPSP is associated with an increased dendritic excitability

To investigate the nature of the fEPSP transformation, we recorded spontaneous excitatory synaptic currents in layer 5 pyramidal cells in patch-clamp mode, in the presence of tetrodotoxin, (mEPSCs – considered to be monoquantal synaptic events (Katz, 1966)). Interestingly, there was no difference in mEPSC amplitude, frequency or kinetics between cells recorded after seizure-like activity was established (Fig S6), compared to the baseline measurements, indicating that the fEPSP transformation does not arise from a sudden potentiation of glutamatergic transmission. Rather, the all-or-nothing nature of the fEPSP transformation, and the increased fEPSP duration (mean 91.75 ±11.87ms), are both strongly suggestive of a postsynaptic effect, in which the optogenetic stimulation evokes dendritic plateau potentials. To test this hypothesis, we imaged Ca^2+^ dynamics in pyramidal apical dendrites in response to stimulation, over the time course of the evolving epileptiform activity (Fig 2, S7, Movies S1-3). Prior to the fEPSP transformation, we found highly localized, low amplitude Ca^2+^ transients limited to the distal tuft. In contrast, after the transformation, each fEPSP was associated with extensive and sustained Ca^2+^ entry throughout the apical dendrites (Fig 2Bi). Furthermore, the transformation in the fEPSP occurred coincidental with a large increase in the Ca^2+^ signal in the apical trunk and somata of layer 5 pyramidal cells, in response to the stimulation (Fig 2Bii, n = 4 brain slices).

**Fig. 2.**
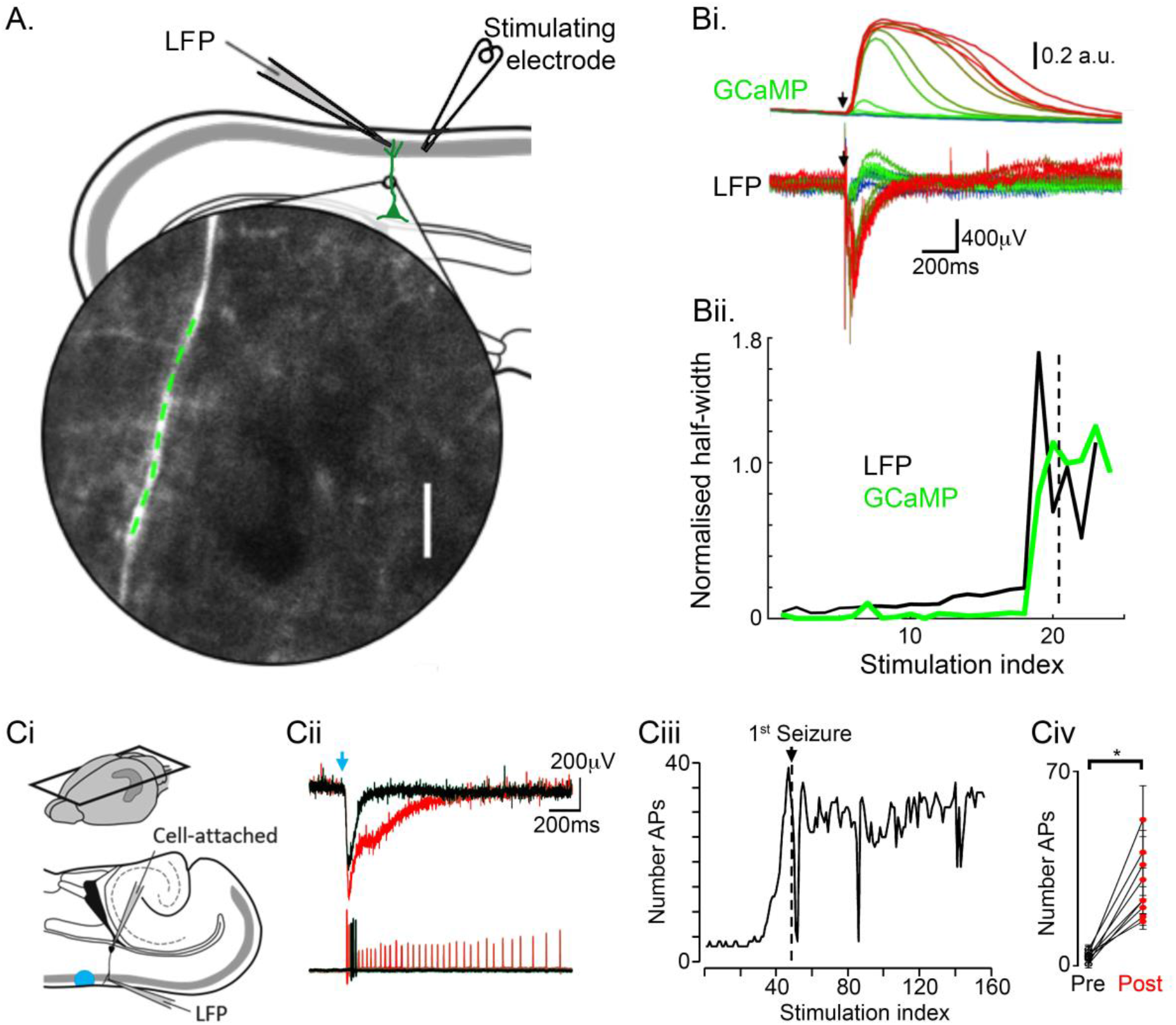
The fEPSP transformation reflects the occurrence of dendritic spiking. (A) Schematic showing the *in vitro* recording arrangement in horizontal brain slices prepared from young adult mice expressing GCaMP6f in pyramidal cells. The inset shows the apical dendrite trunk of a layer 5 pyramidal cell. (Bi) The change in Ca^2+^ signal in the apical dendrite trunk, recorded for sequential stimuli, and the LFP, recorded from a close extracellular electrode. Synchronous events are colour-coded, with green events prior to the transformation, and red events after. (Bii) Plot showing the parallel changes in half-width of both the dendritic GCaMP6f signal (green) and the LFP, for sequential stimuli. (Ci) Schematic showing the recording arrangement for the cell-attached recordings from layer 5 pyramidal cells, and the LFP in the supragranular layers (Cii). (Ciii) Example of the parallel change in the LFP and the spiking pattern of the pyramidal cell, just prior to the first seizure, and (Civ) pooled data showing the large increase in firing in all recorded neurons (n = 9 brain slices, 4 mice; *, p <, paired t- test).

Dendritic plateau potentials in cortical pyramidal cells have previously been noted to be associated with bursts of action potential firing (Larkum et al., 1999). We reasoned that this would constitute a secondary amplification of the optogenetic response, additional to the dendritic potential. We therefore examined the firing patterns of layer 5 pyramidal cells in response to the optogenetic stimulation, during evolving epileptiform activity, using cell attached patch recordings (Fig 2C). In all cases, the fEPSP transformation was associated with a large increase in the number of action potentials per burst (Fig 2Civ; n = 9 brain slices, p = 0.0003, paired t-test). Since this activity delivers excitatory drive back into the local network, all these associated changes-increased dendritic excitability, the consequent increase in dendritic spike probability and burst firing – act synergistically, as positive feedback loops, driving the change from one network state to another.

Together, these experiments indicate that the transformation of the fEPSP represents a shift in dendritic excitability such that a stimulus, which previously had been subthreshold for generating a dendritic spike, suddenly becomes suprathreshold. We therefore examined the cellular processes that underlay this change in dendritic excitability.

## Factors underlying change in dendritic excitability

Since the change in dendritic excitability appeared not to arise from any change in glutamatergic synaptic function, we next asked if it might be explained by changes in inhibitory control(Parrish et al., 2018; Wong and Prince, 1979). We examined the postsynaptic effects of activation of the two main classes of cortical interneuron that target pyramidal neurons. We recorded postsynaptic inhibitory postsynaptic currents (IPSCs) in layer 5 pyramidal cells, in brain slices bathed in 0 Mg^2+^ aCSF, while delivering intermittent (10s period) optogenetic stimulation either to the local parvalbumin-(PV), or somatostatin- (Sst) expressing, interneuronal populations. In both cases, there was a pronounced reduction in the amplitude of optogenetic IPSC during the seizure-like activity, but this only occurred at the onset of the seizure-like event, and only transiently during episodes of intense interneuronal discharges, beforehand (e.g. Fig S8). Furthermore, for both PV and Sst interneurons, the reduction in amplitude recovered fully (Fig 3A; Fig S5) after the ictal event, unlike the fEPSP transformation, which persisted. There are thus two critical differences in the relative timing of the changes in IPSCs and the fEPSP, and so we concluded that the change in the latter does not arise because of altered PV or Sst interneuron function.

**Fig. 3.**
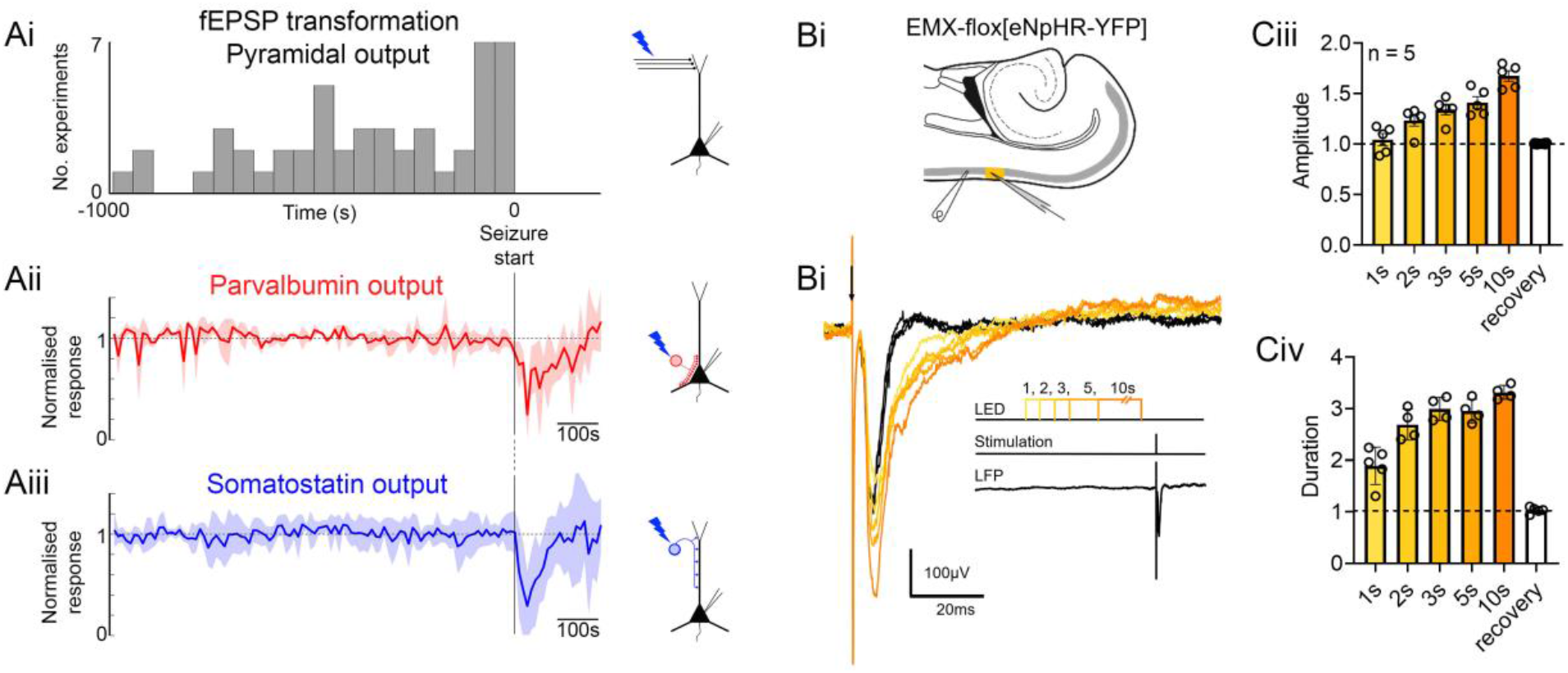
Analysis of inhibitory function, relative to the timing of fEPSP transformation. (Ai) Data from Fig 1F, re-plotted as a frequency histogram, with respect to the timing of the first seizure. On the same time scale, below are shown the amplitude of evoked IPSCs recorded in pyramidal neurons, following optogenetic stimulation of either (Aii) parvalbumin-expressing interneurons (n = 3) or (Aiii) somatostatin-expressing interneurons (n = 7). Example recordings are shown in Fig S5. We only include data for which the recordings were stable for more than 500s prior to and 100s after the onset of the first SLE. For purpose of pooling data from different recordings, each was normalized to the mean over the entire pre-ictal period. Note that the transformation of the fEPSP invariably occurs ahead of the first SLE, but the change in the input of both interneurons to pyramidal cells only occurs at the onset of the SLE, and recovers at its end. (B) Chloride-loading into pyramidal cells, induced by prior activation of the optogenetic chloride-pump, halorhodopsin, is associated with a marked increase in the (Biii) amplitude and (Biv) half-width of the evoked fEPSP.

While synaptic IPSCs appear to be relatively stable in the pre-ictal period, inhibitory function may be compromised in a different way, through rising chloride levels in the postsynaptic neuron. Chloride has been demonstrated to rise sharply at the onset of seizures, both *in vitro*(Ellender et al., 2014) and *in vivo*(Sulis Sato et al., 2017), and the loading continues to progress with subsequent events(Dzhala et al., 2010). Importantly for our observation, chloride levels may rise even prior to the initial seizure-like events, as has been observed both using imaging (Dzhala et al., 2010) and also using gramicidin perforated patch recordings of pyramidal cells, in brain slices bathed in 0Mg^2+^ (Burman et al., 2019). As with the other measures of synaptic function, the time course of the chloride changes does not exactly match the sudden all-or-nothing change in the evoked EPSP, but the fact that there is a progressive creeping rise prior to the onset of seizures suggests that intracellular Cl^-^ levels may still be a critical factor. To investigate this hypothesis, therefore, we first tested whether loading chloride into neurons could lead to the all-or-nothing transformation in the fEPSP. To do this, we used the light-activated chloride pump Halorhodopsin to drive chloride into the pyramidal neuron population, as demonstrated previously (Alfonsa et al., 2015; Raimondo et al., 2012). This reliably induced a large increase in both the amplitude and time course of the fEPSP evoked by electrical stimulation (Figs 3B, S9), supporting the hypothesis that a positive shift in E_GABA_ underlies the fEPSP transformation and associated ictogenesis.

To explore this further, we created an anatomically realistic compartmental model of a pyramidal cell, including two important active conductances, VGCCs and NMDA-receptors (NMDA-Rs) within the dendritic tree, in order to map out the relationship between local E_GABA_ and dendritic excitability. We simulated a transient glutamatergic drive onto the apical tuft, to mimic the experimental optogenetic stimuli, and varied the intensity of stimulation to map out the threshold for a dendritic plateau potential (Fig 4A). The model also included a feedforward inhibition, provided by concomitant activation of a distributed GABAergic synaptic population. We then explored how the threshold for dendritic plateau potentials changed as a function of E_GABA_. We found that, for a given level of excitatory drive, there was always a sudden step increase in the amplitude and duration of the synaptic response, as E_GABA_ becomes more positive (Fig 4B,C). Notably, the threshold glutamatergic drive to trigger a plateau potential, in this model, showed an almost linear inverse relation to E_GABA_ (Fig 4D).

**Fig. 4.**
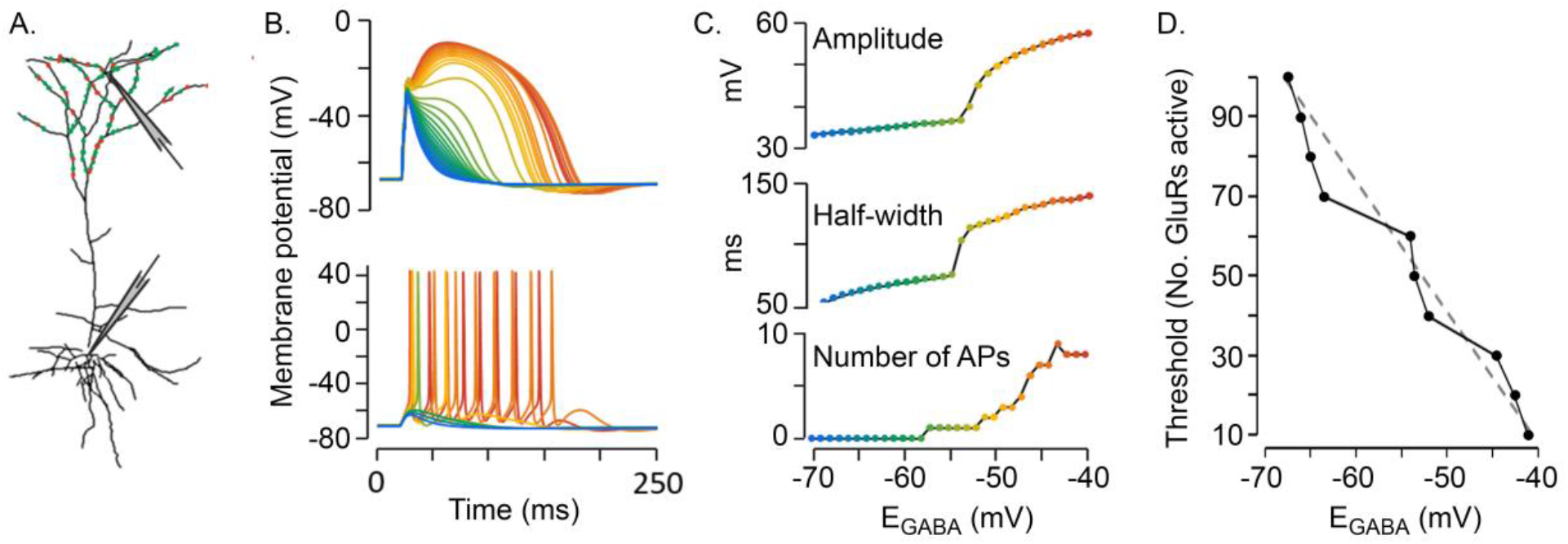
Computational simulations indicate how the threshold for triggering dendritic spikes is inversely related to the GABAergic reversal potential (E_**GABA**_). (A) Structure of the model pyramidal cell, with the locations of the excitatory (green spots) and inhibitory feedforward synapses on the apical tuft. (B) Membrane potential recorded at two sites, in the apical dendrites (top) or at the soma (bottom), showing repeat simulations performed with a constant level of synaptic drive, but with different simulated E_GABA_, colour-coded as shown in (C), which depicts the increasing amplitude and half-width of the plateau potential, and the number of action potentials for each event. Note the step increase for each parameter, as E_GABA_ becomes more positive. (D) Simulations show that the number of glutamatergic synapses required to trigger a dendritic plateau potential is inversely related to E_GABA_.

## Positive feedback mechanisms drive seizure initiation

The occurrence of plateau potentials in pyramidal cell dendrites, and the associated switch to bursting firing, are driven by non-linear (active) conductances, which represent a critical positive feedback mechanism that underlies a rapid escalation in excitability. We link this also to a rise in dendritic [Cl^-^]_i_, which also constitutes a positive feedback: increased network activity leads to increased [Cl^-^]_i_, a consequent reduced inhibitory efficacy, and thus increased network excitability. Raised extracellular [K^+^] can be considered similarly – higher levels of network activity leads to increased [K^+^]_o_ which in turn increases excitability – and, notably, is linked also to chloride levels by the cation-chloride co-transporters, meaning that raised [K^+^]_o_ also drives raised [Cl^-^]_i_. Dendritic potentials obviously occur during normal cortical function, but the risk of a seizure escalates if these are coordinated in many neurons simultaneously. Raised extracellular K^+^ is one mechanism that might create such a coordinated state, although by no means the only one. Others may include alterations in the expression of various ion channels in neuronal dendrites(Hoffman et al., 1997; Lerche et al., 2013; Shah et al., 2004), or a change in the functionality of Somatostatin (Sst) interneurons. While most attention has focused upon the role of parvalbumin (PV) in opposing epileptic discharges, in fact both sets of interneurons are very active immediately ahead of an ictal wavefront(Parrish et al., 2019), and dysfunction of either class can lower the threshold for seizures(Calin et al., 2018; Khoshkhoo et al., 2017; Sessolo et al., 2015; Wang et al., 2020).

Increases in intracellular [Cl^-^] and extracellular [K^+^], have both been identified previously in many studies as playing a role in ictogenesis, although their relative importance continues to be debated. Both are difficult to assay using conventional passive recording of brain activity, but the simple stimulation paradigm we use here provides an indirect means of monitoring them, by assaying, instead, the threshold for a dendritic plateau potential. In our experimental models, this optogenetic stimulation provides accurate advanced warning of the first seizure, to within a few minutes. Furthermore, monitoring seizure risk in this way, using an active probing between 0.1- 0.033Hz, has itself an anti-epileptic effect, thus constituting a “double-win”.

The interrelated nature of these various positive feedback mechanisms provide a coherent explanation for the suddenness with which a seizure-prone brain switches from a normal physiological state into a seizure state: together, they create an unstable point at what might be termed the seizure threshold, the bifurcation point for the network. Once this is surpassed, the functional state of the network rapidly accelerates towards the seizure state. For expediency of experimentation, we utilized mainly acute in vitro models, begging the question whether the same progression applies also in chronic epilepsy. Together these provide experimental support, and a cellular mechanistic explanation, for the theoretical exploration of attractor models applied to epileptic transitions at the network level(Chang et al., 2018b; Jirsa et al., 2014). Recent research into the cellular mechanisms of ictogenesis has focused predominantly, in recent years, onto inhibitory dysfunction, but our study redirects our attention towards the conductances that underlie plateau potentials(Johnston and Brown, 1981; Traub and Miles, 1991; Traub et al., 1989; Wong and Prince, 1979). By linking these various cellular phenomena, we draw together several major lines of epilepsy research.

## Materials and Methods

### Ethical Approval

All animal handling and experimentation were done according to the guidelines laid by the UK Home Office and Animals (Scientific Procedures) Act 1986 and approved by the Newcastle University Animal Welfare and Ethical Review Body (AWERB # 545). For selective expression of channelrhodopsin2 (ChR2), halorhodopsin (HR), or GCaMP6f under neuronal subtype-specific promoters EMX-1 (B6.129S2-Emx1tm1(cre)Krj/ J; Jackson Laboratory stock number 5628), PV (B6;129P2-Pvalb < tm1(cre)Arbr>/J; Jackson Laboratory stock number 8069), or Sst (B6N.Cg.Ssttm2.1 (cre)Zjh/J; Jackson Laboratory stock number 18973), animals expressing the floxed gene of interest were crossed with animals expressing Cre under the appropriate promoter. Widespread cortical expression of ClopHensor was reliably induced following the injection of EMX-1-Cre mice at postnatal day 0-1 with flox[EF1α-ClopHensor] in an AAV8 viral vector.

### Slice preparation

Young adult C57BL/6 mice (age 6-24 weeks) were used in this study. Mice were housed on a 12h light/dark cycle, and were given access to food and water *ad libitum*. Mice were sacrificed by cervical dislocation, the brain removed, and sliced in ice cold cutting solution (in mM): 3 MgCl2; 126 NaCl; 26 NaHCO3; 3.5 KCl; 1.26 NaH2PO4; 10 glucose. Horizontal slices containing both neocortex and hippocampus proper (z = −3.1–4.7mm) were taken using a Leica VT1200 vibratome (Nussloch, Germany) at a thickness of 300µm for patch recordings, and 400µm for field recordings. Slices were immediately transferred to a holding chamber and incubated at room temperature for >45 min prior to recording in artificial cerebrospinal fluid (aCSF) containing (in mM): 2 CaCl_2_; 1 MgCl_2_; 126 NaCl; 26 NaHCO_3_; 3.5 KCl; 1.26 NaH_2_PO_4_; 10 glucose. All solutions were bubbled with carbogen (95% O_2_, 5% CO_2_) throughout.

### *In vitro* electrophysiology

Local field potentials were recorded in interface recording chambers in which slices were superfused with carbogenated aCSF maintained at a temperature of 34±1°C. The rate of the perfusate circulation was maintained with a peristaltic pump at ∼3ml min^-1^ (Watson Marlow). Borosilicate glass microelectrodes (GC120TF-10; Harvard apparatus, Kent) were pulled using an electrode puller (Model-P87, Sutter Instruments, CA, USA) and filled with aCSF. Recordings were obtained with electrodes at a resistance of 1-3MΩ. Analog signals were acquired using in-house built headstages (10x gain) connected to BMA-931 AC (0.1Hz) differential amplifier (Dataq instruments, Akron, USA) with the gain set appropriately for the recording between 200-500. Amplified signals were digitised at ≥10kHz using a Micro 1401-3 data acquisition unit (Cambridge Electronic Design, UK), bandpass filtered at 1-3000Hz, and stored on a computer using Spike2 (V7.2 or later) acquisition software (Cambridge Electronic Design, UK).

For patch clamp recordings, slices were bathed in carbogenated aCSF perfused at 3-5 ml min^-1^ and heated to 34±1°C. Recordings were made using 4-7MΩ borosilicate glass microelectrodes (GC150F-10, Harvard apparatus, Kent). Electrodes were filled, unless otherwise stated, with K^+^-gluconate-based filling solution containing (in Mm): 125 K-gluconate, 6 NaCl, 10 HEPES, 2.5 Mg-ATP, 0.3 Na_2_-GTP. Electrode filling solutions were pH adjusted to 7.3 with KOH and osmolarity adjusted to 280-290mOsm. Patch clamp data were acquired using pClamp software v10.5, Multiclamp 700B, and Digidata acquisition board (Molecular Devices, CA, USA). Signals were digitised with a sampling frequency of 10 kHz.

Two *in vitro* epilepsy models were used: the zero-magnesium (0Mg^2+^) and 4-AP models. In both cases, baseline recordings were established in normal bathing media initially. Then, we either washed out Mg^2+^ ions or added the potassium channel blocker, 4-aminopyridine (100µM). For halorhodopsin chloride loading experiments, slices were prepared from mice bred to express enhanced Halorhodopsin (eNpHR3.0) in pyramidal cells, under the CamK2a promoter (cross bred from two Jackson laboratory lines: #014539 and #5359). Extracellular recordings were made from the supragranular layers of cortex, following periods of illumination for several seconds (1-10s) with 561nm yellow light, to activate the Halorhodopsin pump. Similar illumination has been shown by previous work to induce a positive shift in E_GABA_ of up to 20mV (Alfonsa et al. 2015). Each illumination was followed by an electrical stimulation, delivered to superficial neocortex (0.1-4V, 100µs duration) using a Digitimer DS3 electrical stimulator. This stimulation was delivered at a delay of 500ms with respect to the light-off, to allow us to separate the electrographic effects of the optogenetic activation from the evoked field EPSP.

### *In vivo* electrophysiology

For acute *in vivo* recordings, mice were initially anaesthetised by intraperitoneal injection of urethane (20% w/v in 0.9% sterile saline) at a dose of 0.1ml per gram weight. Anaesthesia was supplemented during craniotomy surgery with 1-2% isoflurane (Isothesia, Schein Animal Health) in oxygen 1-1.2L min^-1.^ Mice were head fixed in a stereotactic frame. Isoflurane was discontinued once the craniotomy was complete, and at least 20 minutes prior to recording. Internal temperature was monitored by a rectal thermometer and maintained with a thermoregulatory blanket. A dental drill (RAMPower, RAM) was used to make small craniotomies (typically 2-3mm diameter), the dura removed to facilitate penetration with the multielectrode probes, and the cortical surface kept moist with 0.9% saline. Electrophysiological signals were recorded using a 4×4 multielectrode array (NeuroNexus, electrode separation 200μm, recording sites 1250μm^2^, impedance <3MΩ at 10kHz. Signals were amplified and digitised by a Plexon AC amplifier (MAP Data Acquisition, Hkl3, PLEXON), and were amplified and digitised at 25kHz. Data was stored using Sortclient (V3) software. Optogenetic stimulation was delivered from an adjustable-intensity mountable 470nm LED, through an optic fibre and matched cannula (Ø400μm, ThorLabs) driven from Spike2 via a Micro 1401-3 data acquisition unit (Cambridge Electronic Design, UK).

### Calcium imaging

Brain slices at a thickness of 400μm were submerged in aCSF and perfused at a rate of 3-4ml min^-1^ and heated to 34±1°C. Local field potentials were recorded with extracellular electrodes described above, and digitised at 10kHz (Digidata 1440, Multiclamp, Axon Instruments), and stored using Thorlabs software (ThorSync, V3.2, and ThorImage, V3.2). Images were captured using a Bergamo II 2-photon microscope (ThorLabs), exciting the sensor with a MaiTai laser (SpectraPhysics) at 940nm through a 16X objective lens (water immersion, NA 0.8, Nikon). Acquisitions were taken at 30Hz in 512×512 resolution, scanning in both directions with a galvo-resonant scan head. 1.5s imaging periods were centred around electrical stimulations (0.1-4V, duration of 100μs, Digitimer DS3) delivered at 0.016Hz (stimulating every 60s), via chlorinated silver wires in a θ-glass borosilicate electrode filled with aCSF, for the duration of the experiment.

### Computational modelling

A compartmentalised single-cell model was developed using the NEURON simulation environment (Hines and Carnevale, 2001). The 3D morphology of the model was adapted from Louth et al. (Louth et al., 2018), for investigating the effects of changing E_GABA_ on the response to synaptic excitation in the apical tuft. As values for channel distribution and conductance in cortical pyramidal cells have previously been empirically measured and published for the NEURON modelling environment, published values were used to constrain this model. Typical values for the biophysical properties of the cell were used, with Ri = 150 Ω cm^-1^, resting Rm = 60 kΩ cm^-2^, and Cm = 1 µF cm^-2^ (Migliore et al., 2003), and were set to be uniform across the cell. I_h_(0.0002 mS cm^-2^) and leak currents were included with reversal potentials set to −30mV and −90mV respectively (Migliore et al., 2003). Classic Hodgkin-Huxley conductances were included at the soma; KV (type A) channels at conductance of 0.04mS cm^-2^, NaV channels at 0.0005 mS cm^-2^. Excitatory synapses were modelled using an extension of the Exp2Syn class in NEURON. Briefly, the Exp2Syn class is a general class for synapses with a conductance described by a sum of two exponentials with rate constants τ_rise_and τ_decay_, here denoted τ_1_ and τ_1_.

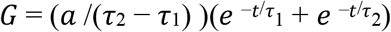

In excitatory synapses the values for time constants for AMPA (τ_rise_ = 0.2ms; τ_decay_ = 2ms) and NMDA receptors (τ_rise_ = 3ms; τ_decay_ = 35ms) were derived from the empirical measurements made by Shulz et al (Schulz et al., 2018). The threshold for voltage-dependent unblocking of NMDARs was set at-16mV (Kampa et al., 2004). The conductances for both AMPA and NMDA were set to 0. 14 nS, at synapses at a density of 0.5 µm^-2^ (Bloss et al., 2016; Kampa et al., 2004). The voltage dependency of the NMDAR current was modelled as in Shulz et al. (Schulz et al., 2018) with the function:

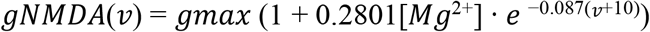

Where *v* is in mV and the Mg^2+^ concentration is 1mM.

At inhibitory synapses both rectifying and non-rectifying GABA_A_ receptors were included, as outward rectifying channels were reported to powerfully constrain dendritic depolarisation (Schulz, et al., 2018). GABA_A_ synapses were included with a conductance of 0.7 nS, with τ_rise_ = 0.5ms, and τ_decay_ = 15ms (Schulz et al., 2018). Outward rectifying GABA_A_ channels were again modelled by expanding the Exp2Syn class (Schulz et al., 2018), and τ_rise_ to 1ms, and τ_decay_ to 30ms, for membrane voltages above −52mV. Subsets of GABAergic synapses at 0.1 µm^-2^ were activated simultaneously with excitatory synapses. Synapse numbers and distribution were taken from Bloss et al (Bloss et al., 2016). Voltage-gated calcium channel (VGCC) conductances were included as described in Lazarewicz et al. (Lazarewicz et al., 2002), which implements L- and T-type calcium channels according to their distribution in CA1 pyramidal neurons. Here Ca_L_ and Ca_T_ conductances have been added. The density of Ca_L_ conductance is set across the dendritic tree at 0.0013 mS cm^-2^, and Ca_T_ in the soma and dendrites within 100µm path distance to the soma at 0.001 mS cm^-2^ (Lazarewicz et al., 2002). In order to simulate lateral cortical afferent stimulation, and replicate the experimental protocol, only the apical tuft was stimulated. Excitatory and inhibitory synapses were activated simultaneously to simulate feedforward inhibition which is typically concomitant with excitatory drive *in vivo* (Buzsaki et al., 2007; Doron et al., 2017). Increasing numbers of excitatory synapses in the dendritic tuft were activated for a range of values of E_GABA_ (−70 to −40mV) to identify the threshold level of excitation for eliciting a plateau potential. No proximal synapses were added, and no background synaptic noise was included, for simplicity. For clarity of presentation, somatic Na_V_ channels were silenced to generate the dendritic potentials presented.

### Data analysis

Data were analysed by custom MATLAB code, and automated wherever possible.

## Acknowledgments

We would like to thank all members of the Trevelyan laboratory for discussion and help throughout the project. We also thank the support staff in the animal facility at Newcastle University. We also thank Dr Sasha Gartside for help establishing the *in vivo* recordings, Dr Frances Hutchings, for help with the NEURON simulations, and Dr Yujiang Wang, Chris Papasavvas, and Professors Gimmi Ratto, Massimo Scanziani and Mike Hausser for comments on the manuscript.

## Funding

This work was supported by a Newcastle University PhD studentship (RG), and grants from MRC, Epilepsy Research UK, Wellcome Trust and EPSRC.

## Author contributions

Design of the project, AT, RG; Experimental recordings, RG, RP, LA, LO, AT; Analyses, RG, LA, AT; Computer simulations, RG; Figure preparation, RG, AT; Writing manuscript, AT, RG; all authors reviewed the final manuscript.

## Competing interests

No competing interests

## Data and materials availability

All data is available in the main text or the supplementary materials.

## Supplementary Materials

Figures S1-S9

Movies S1-S3

## Supplementary figures

**Figure S1.**
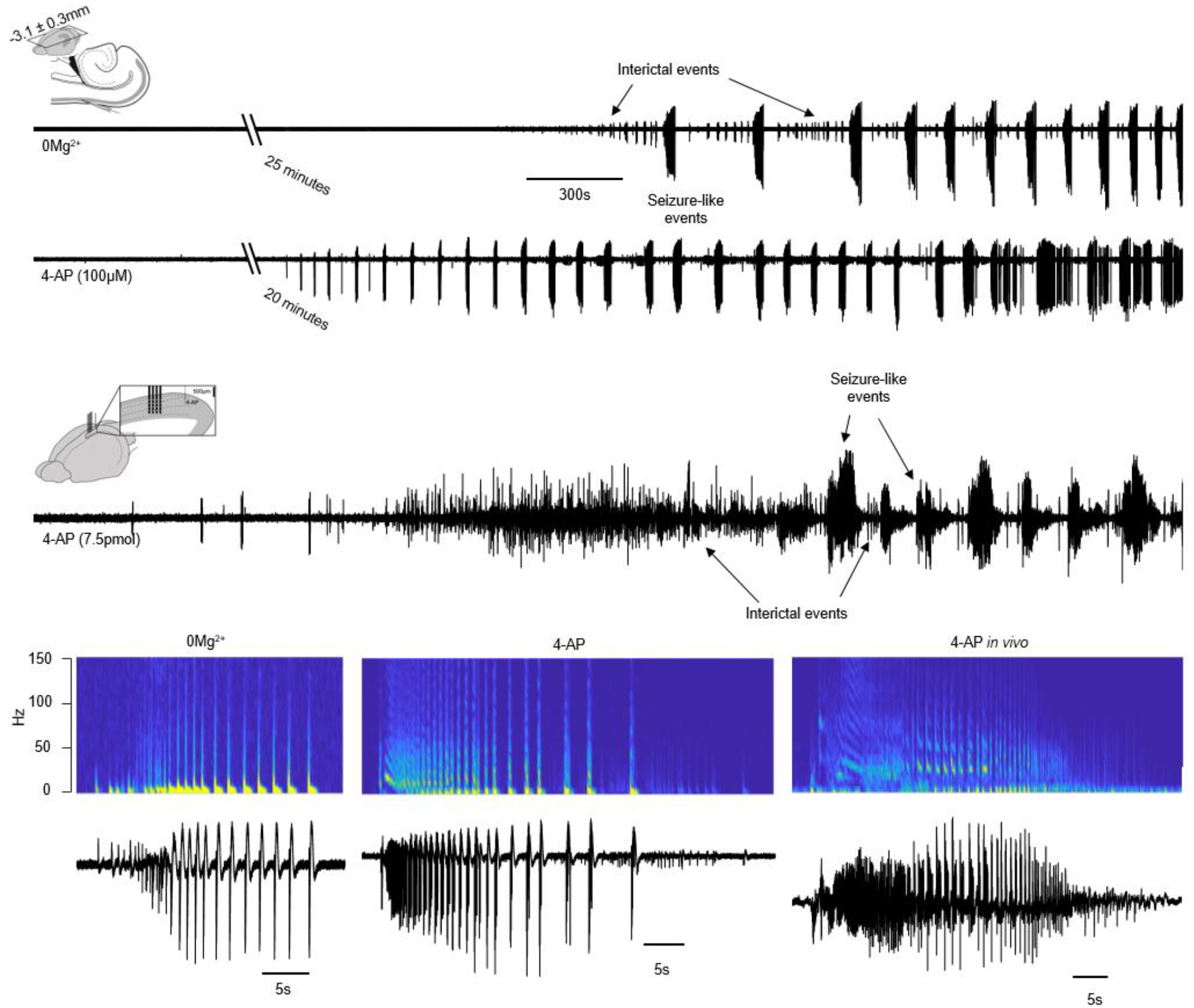
Characteristic development of the acute epilepsy models used (top, 0Mg^2+^ and 4-AP *in vitro*; bottom, intracortical injection of 4-AP *in vivo*). Both models develop with a typical latency after the pharmacological manipulation has been applied which can be used to study seizure susceptibility. After this delay, these models both evolve recurrent seizure-like events with distinctive after-discharges (clear in lower panel). These events can easily be identified, with remarkable similarities in their time-frequency composition, and conserving many properties of electrographically recorded seizures in epileptic patients. Both models also show interictal activity; short burst of hypersynchronous activity which does not evolve into a sustained seizure-like event.

**Figure S2.**
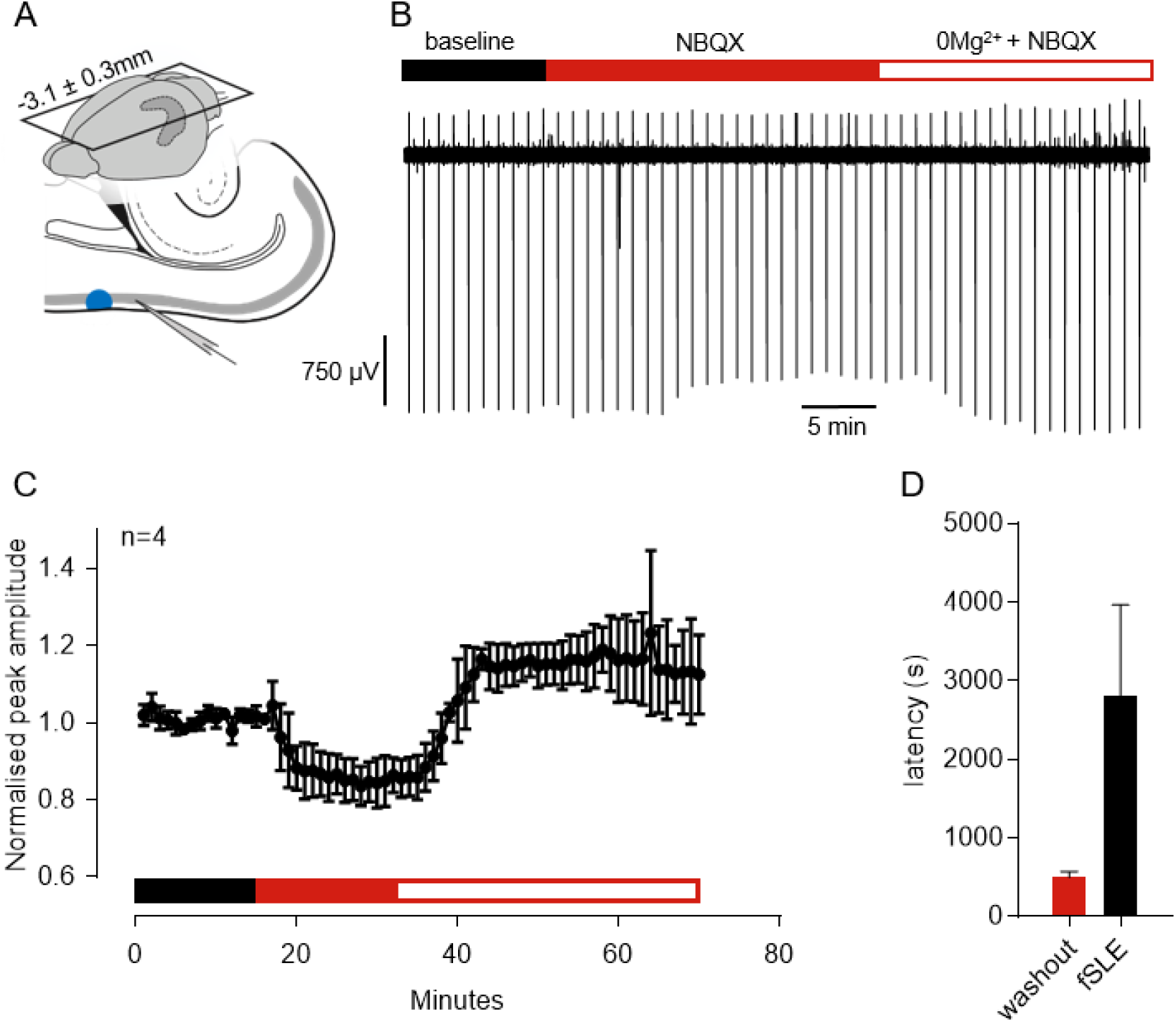
Demonstration of the time-course of the direct effect of the washout of Mg^2+^ from the perfusate. **A**. Schematic of recording strategy in acutely prepared horizontal brain slices containing hippocampus, neocortex, and entorhinal cortex. Blue dot indicates the location of optogenetic illumination with 470nm light. **B**. Representative trace, showing decrease in increase in evoked response amplitude following blockade of AMPA receptors with NBQX, and subsequent washout of magnesium ions from the perfusate. **C**. Grouped slice data showing consistency in both the decrease and increase in response amplitude following manipulations. **D**. Comparison of the time-course of the washout of magnesium, as calculated from the plateau of data shown in C, with the onset latency of seizure-like activity in the 0Mg^2+^ model in horizontal brain slices.

**Figure S3.**
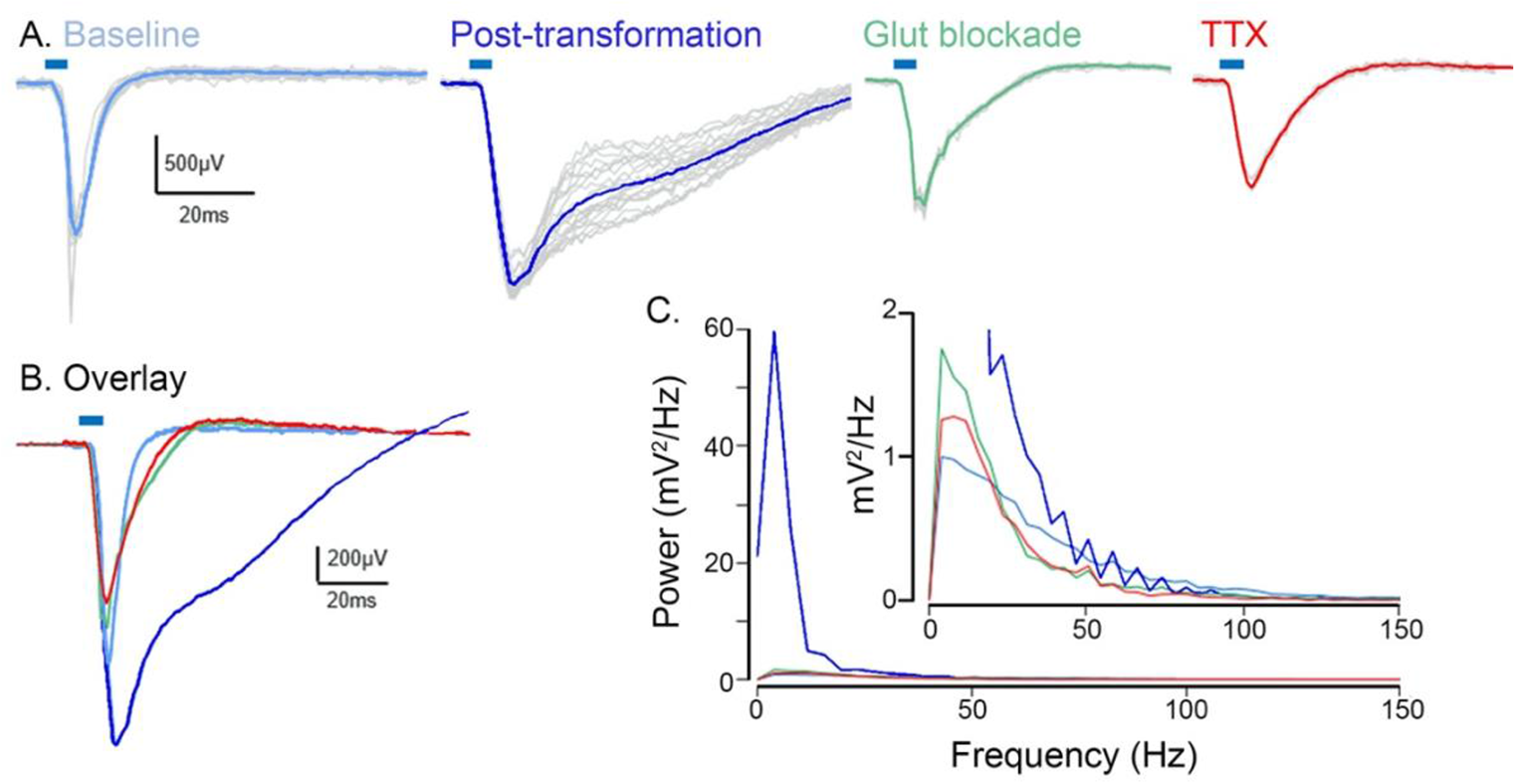
The optogenetically evoked field event can be deconstructed pharmacologically, by applying sequentially glutamatergic blockers (green trace: AP5, 50µM and NBQX, 40µM) and then tetrodotoxin (1µM, red trace). The similarity between the red trace and the initial events, indicates that the baseline evoked responses had a large component that is directly attributable to the channelrhodopsin current.

**Figure S4.**
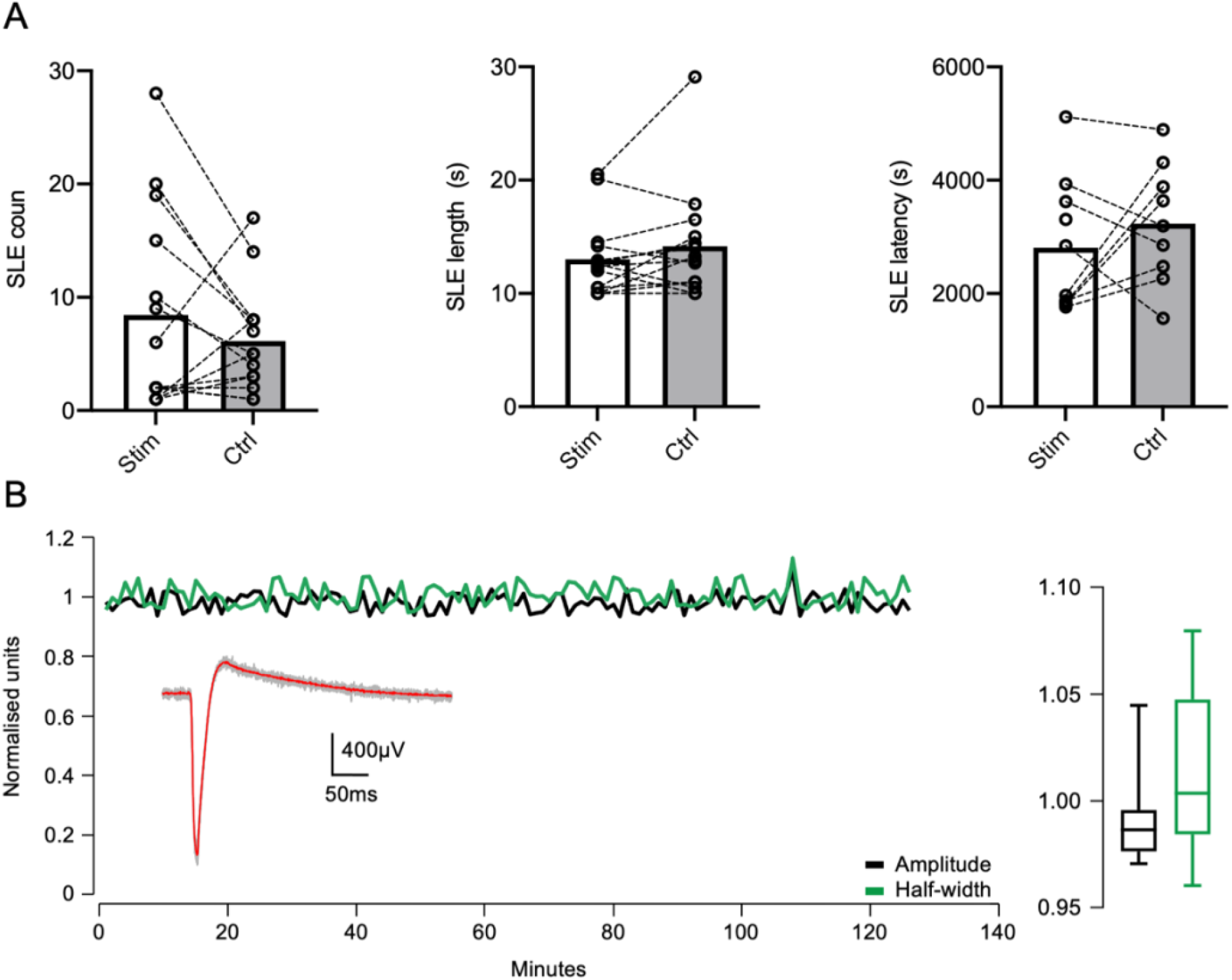
Stimulations at an interval of 60s (0.016 Hz) do not significantly alter the progression of seizure-like activity in the 0Mg^2+^ model, and the evoked responses are stable over hours. **A**. Demonstration that stimulations at an interval of 60s do not alter the onset latency (left), seizure instance (middle), and average seizure length (right), when compared to non-stimulated controls (grey squares). **B**. Representative measures of amplitude and half-width from a single recording, showing the ChR2- evoked responses are stable in baseline conditions on the order of hours. Inset, all responses from the trace overlaid with the averaged response (red). Left, responses do not vary >5% of their mean value over the course of the recording.

**Figure S5.**
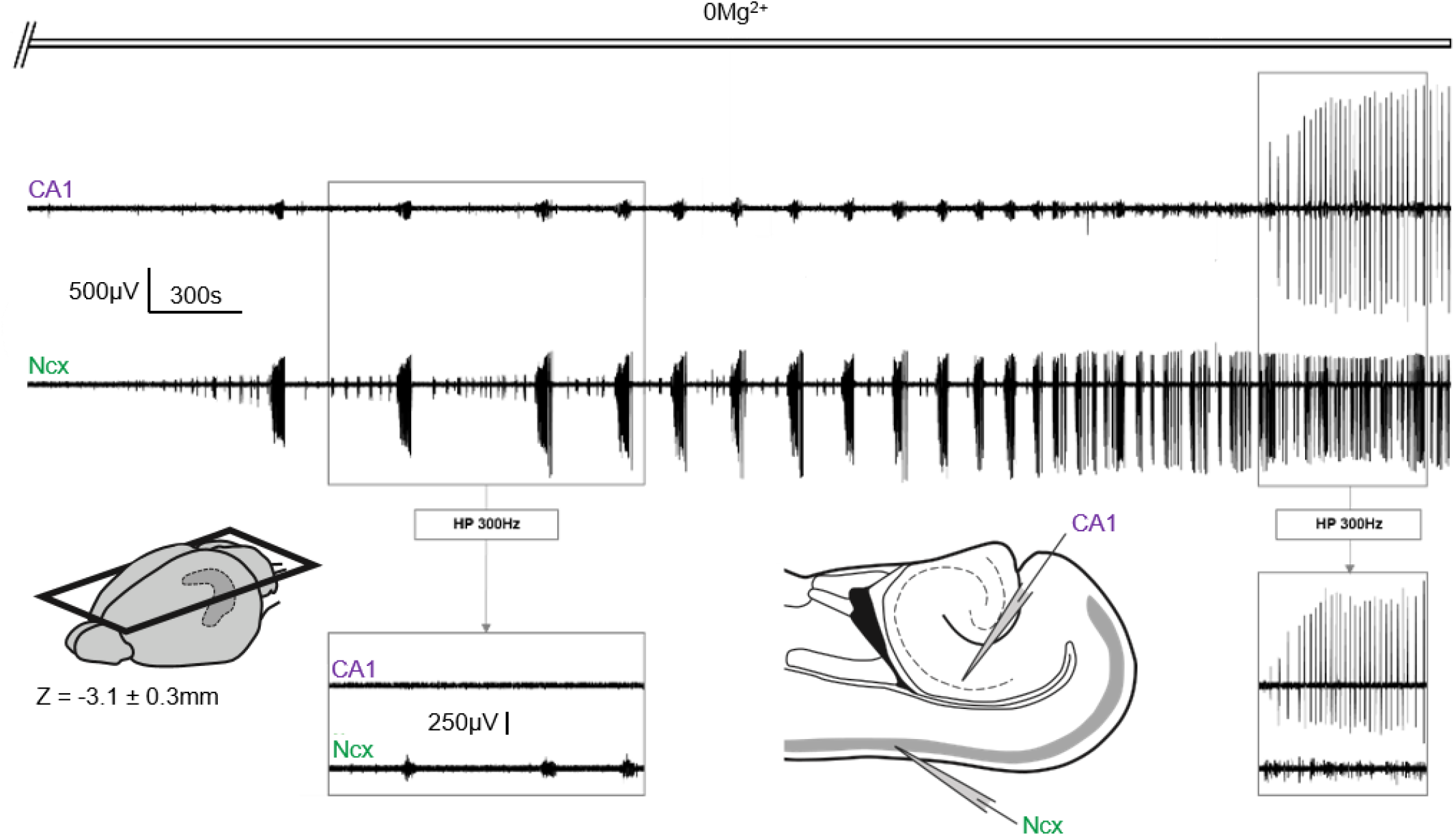
Characteristic development of epileptiform activity in the 0Mg^2+^ model recorded in the supragranular layers of neocortex and *s. pyramidale* CA1 subfield of neocortex. Acutely brain slices were taken horizontally in order to include both regions and preserve entorhinal connections. Ictal activity develops with different latencies in the two regions, first in neocortex, where sustained and recurrent seizure-like events are established. These events show no high-frequency correlate in the hippocampal formation, indicating that the deflections recorded at that site are field effects only from ongoing ictal crises in neocortex (left box). These seizure-like events gradually decrease in interval until the recruitment of hippocampus to the model, at which point spike-wave discharges originating in CA3 synchronise both regions into metronomic discharges (LRD), highlighted in the right box.

**Figure S6.**
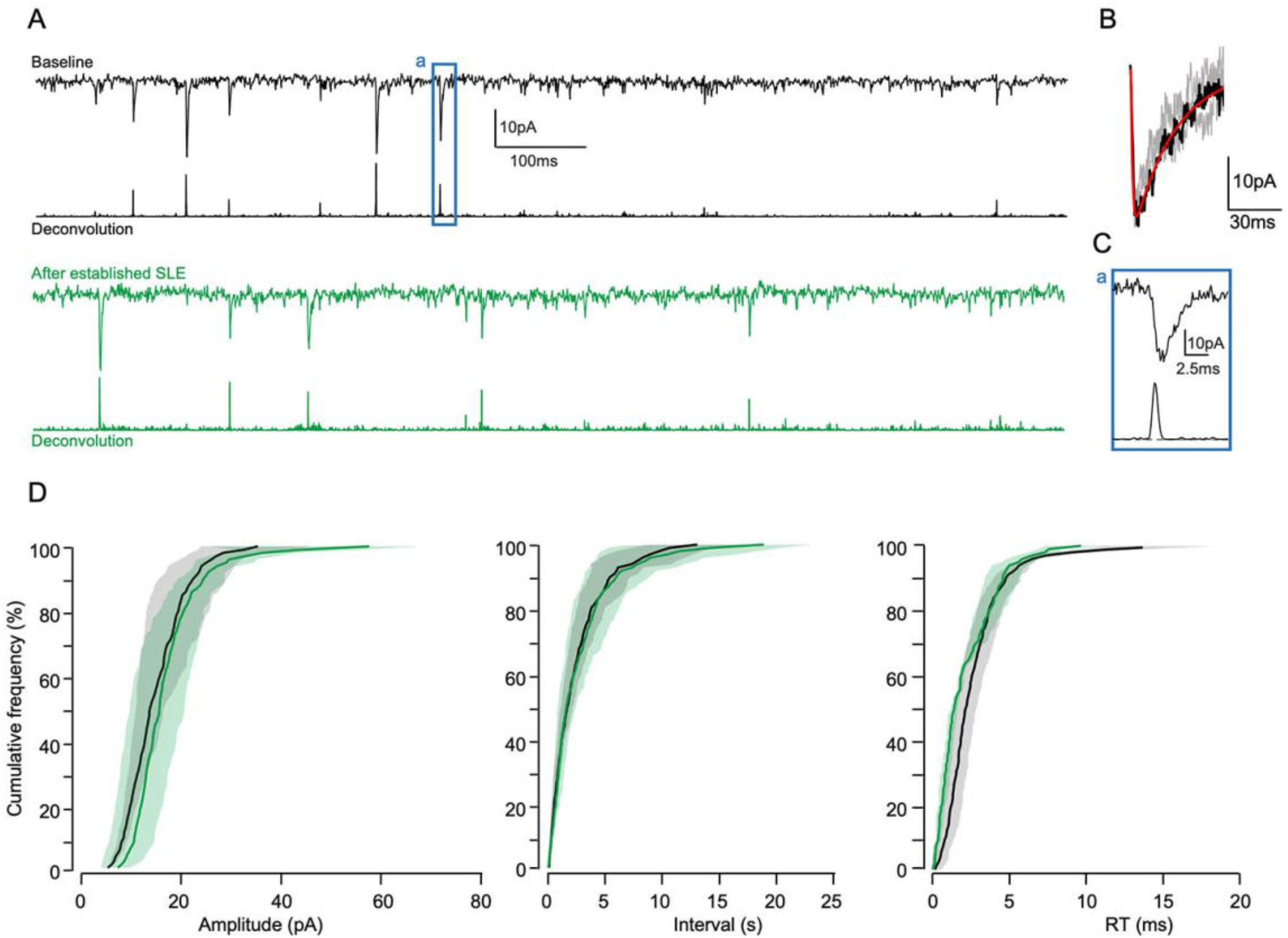
Analysis of miniEPSCs in baseline aCSF and after SLE are established in the 0Mg2model. A. Representative patch-clamp recordings of L5 pyramidal cells in the presence of gabazine and TTX showing spontaneous miniEPSCs. B. Example fitting of sum-of-exponentials curve to manually confirmed miniEPSCs in the trace. This curve is used to produce the deconvolved traces shown in C.C. Deconvolution traces showing how deconvolution of the raw patch-clamp traces in A with the scaled template shown in B provides reliable identification of even small miniEPSCs. D. Zoom of box a in C showing that deconvolution provides excellent identification of miniEPSCs regardless of amplitude. E. Comparison cumulative frequency plots showing (from left to right) amplitude, interval, and rise-time of recorded miniEPSCs, and showing no significant difference between those recorded in baseline conditions and those recorded after the onset of SLE in the presence of 0Mg2+.

**Figure S7.**
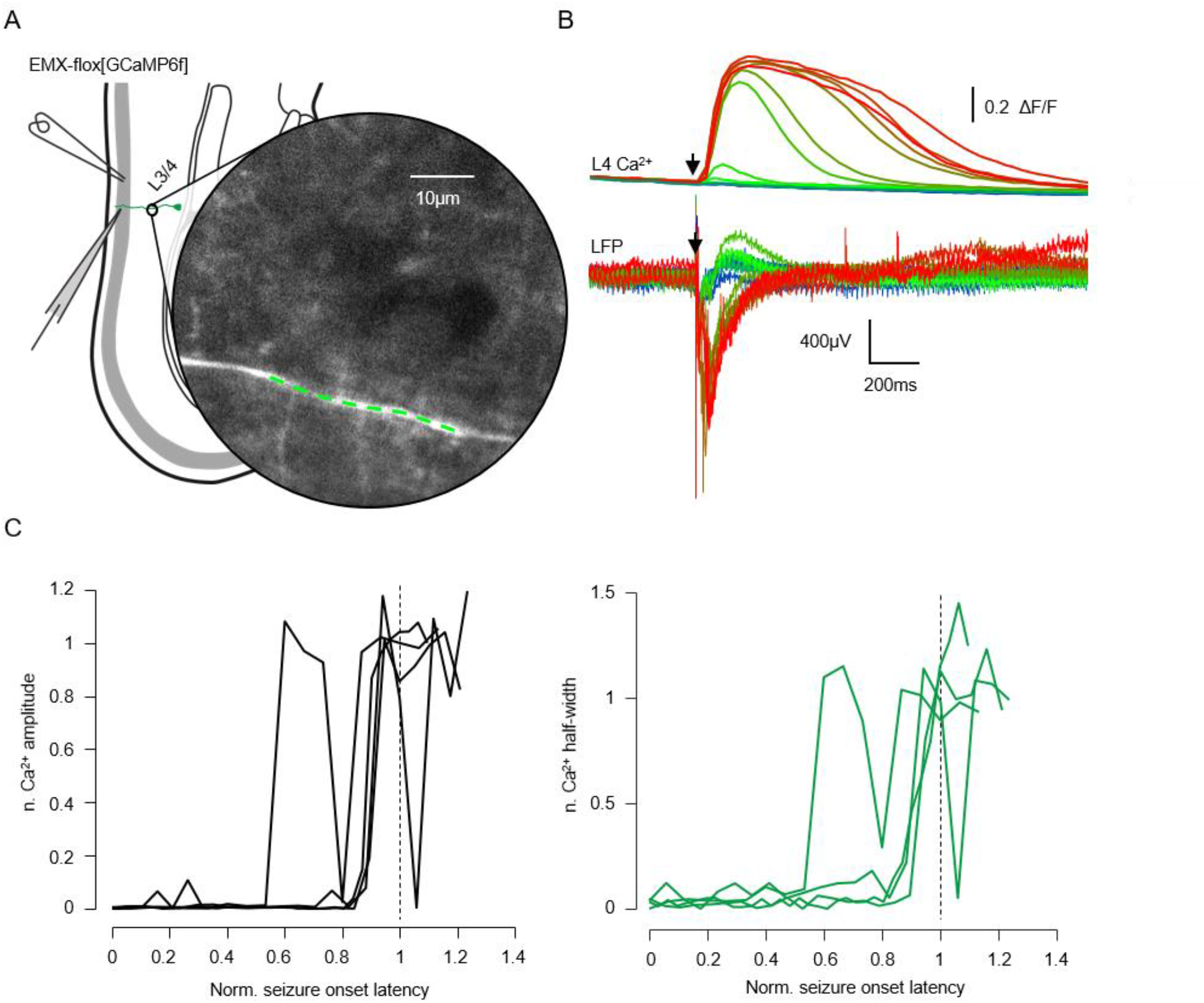
Summary figure showing substantial increase in the amplitude and time-course of calcium transients in the apical dendrite (L4). A. Recording strategy. LFP recorded from L2 as per previous experiments. GCaMP6f was constituently expressed in pyramidal cells, and imaged in the proximal apical dendrite. B. Representative example events showing consistency between the evoked responses in the superficial LFP and calcium transients recorded from the apical dendrite. Black arrow indicates the time of electrical stimulation. C. (Left) Normalised peak amplitude and half-width (right) of calcium transients evoked by electrical stimulation with the time between the application of 0Mg^2+^ aCSF and the onset of the first SLE normalised for direct comparison.

**Figure S8.**
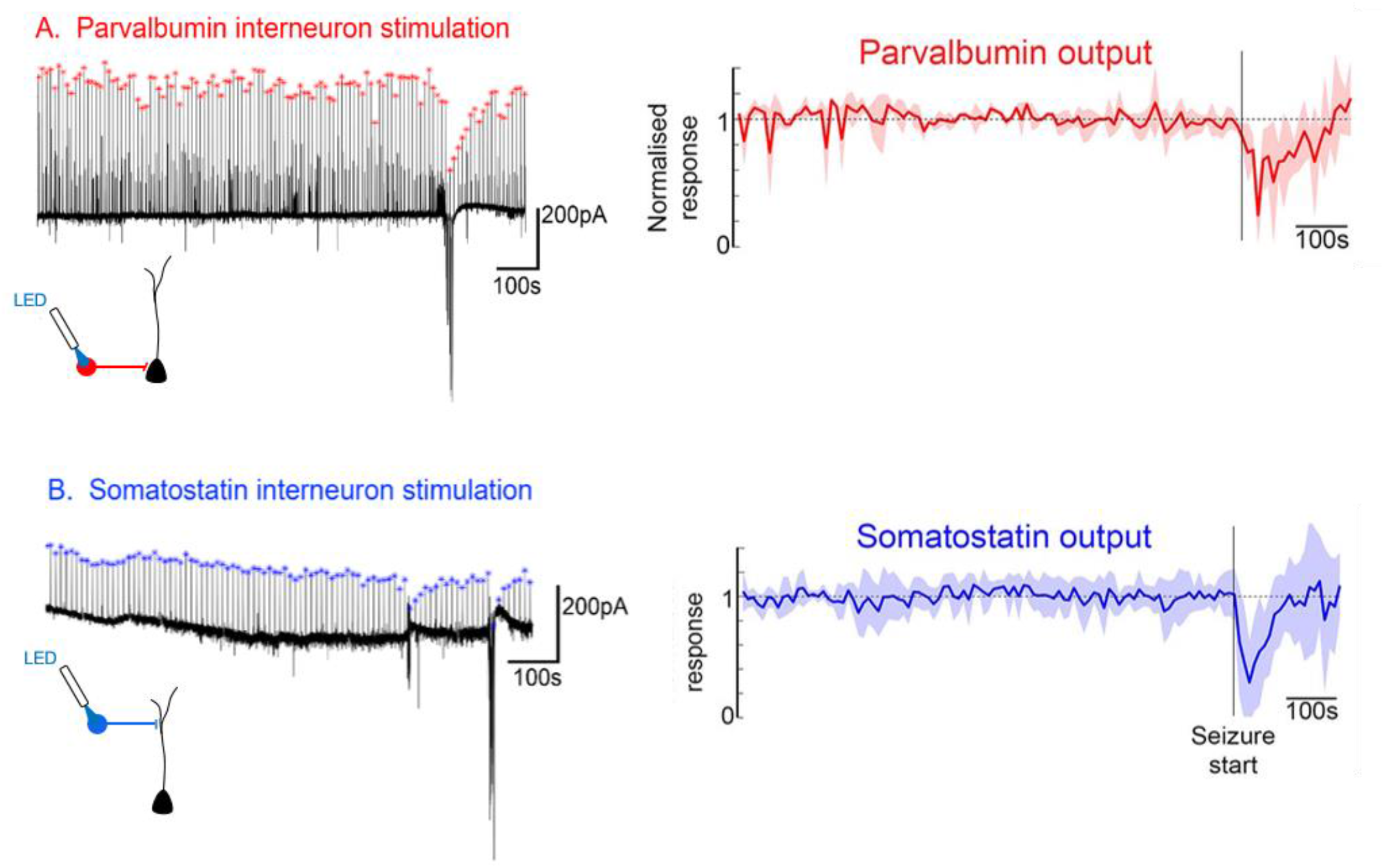
Example voltage clamp recordings of the postsynaptic response, in Layer 5 pyramidal cells, to optogenetic stimulation of either parvalbumin-expressing, or somatostatin-expressing interneurons. Note the transient nature of the drop in the post-synaptic currents at the time of the SLE, or following an intense burst of network activity (see lower recordings ∼200s prior to the SLE). The right panels show the normalized response plots, which are reproduced here from Fig 3 in the main text.

**Figure S9.**
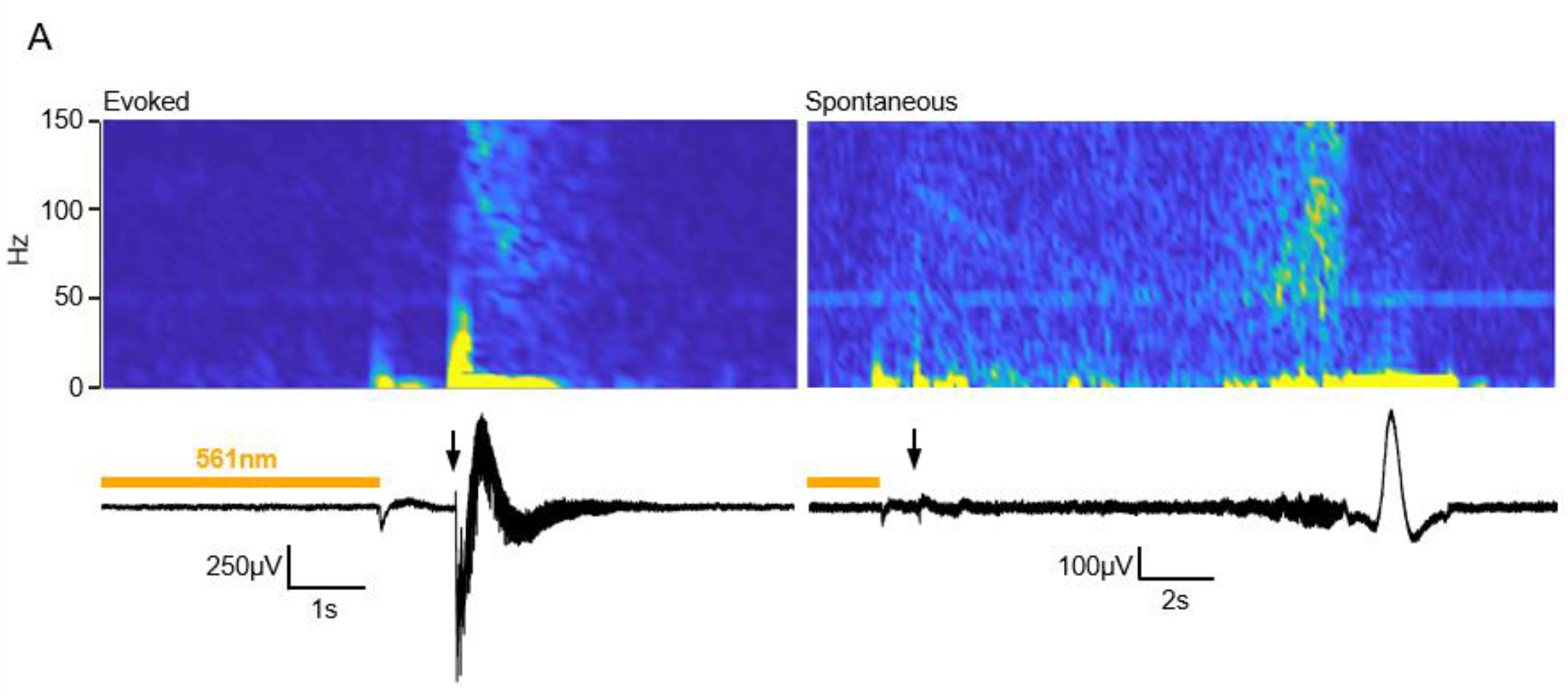
Field recordings of epileptiform discharges occurring in brain slices prepared from non-epileptic mice which are bathed in 1mM (normal) aCSF, but which experienced intermittent trials of intense and prolongued (10s) activation of the optogenetic chloride pump, Halorhodopsin. Without the Halorhodopsin activation, such slices never show such discharges. We showed previously that this pattern of Halorhodopsin activation causes a substantial positive shift in the GABAergic reversal potential (Alfonsa et al, 2015) After a period of chloride-loading induced by Halorhodopsin activation, electrical stimulation induced epileptiform discharge (left panel) on most trials, with very short latency. On other trials, the epileptiform discharge did not occur immediately on stimulation, but instead, the activity built up and a spontaneous event occurred with a delay.

### Movie legends

**Movie S1. Layer 2 pre-transformation**. Movie shows the apical dendrites of pyramidal cells, expressing GCaMP6f under the Emx1 promoter, during a stimulation epoch prior to the transformation of the field EPSP. The movie plays in real time, and the stimulation occurs about half a second into the movie, when one can see a handful of dendritic segments light up. Scale bar = 50µm.

**Movie S2. Layer 5 pre-transformation**. This movie was made using identical microscope settings to Movie S1, but the field of view is approximately in layer 5. Around half a second in, one can see a transient increase in fluorescence in a single dendrite just to the right of the arrows. The movie plays in real time. Scale bar = 50µm.

**Movie S3. Layer 5, post-transformation**. Movie showing the same field of view as Movie S2, and with the same microscopy settings, and the same strength electrical stimulation, but made after the transformation in the field EPSP. Note the very large and widespread Ca^2+^ signal now associated with the stimulation. The movie plays in real time. Scale bar = 50µm.

## Notes

### Competing Interest Statement

The authors have declared no competing interest.

